# Enterovirus 71 structural viral protein 1 promotes mouse Schwann cell autophagy via endoplasmic reticulum stress-mediated peripheral myelin protein 22 upregulation

**DOI:** 10.1101/314468

**Authors:** Pei-qing Li, Si-da Yang, Dan-dan Hu, Dan WeEI, Jing Lu, Huan-ying Zheng, Shu-shan Nie, Guang-ming Liu, Hao-mei Yang

## Abstract

Enterovirus 71 (EV71) accounts for the majority of hand, foot and mouth disease-related deaths due to fatal neurological complications. The clinical observations and animal models found the early invasion of nervous system, and the demyelinating phenomenon was observed. As one of the receptors of EV71 structural viral protein 1 (VP1), SCARB2 mainly exists on the myelin sheath. EV71 VP1 can promote viral replication through inducing autophagy in neuron cells. This study aims to investigate the role and mechanism of VP1 in autophagy of mouse Schwann cells (MSCs). An EV71 VP1-expressing vector (pEGFP-C3-VP1) was generated and transfected into MSCs. Transmission electron microscopy (TEM) and Western blot analysis of the autophagy marker microtubule-associated proteins 1A/1B light chain 3B (LC3B) were used to assess autophagy in the cells. Real-time PCR and immunofluorescent staining were performed to determine the expression of PMP22. Small interfering RNA against PMP22 was employed to investigate the role of PMP22 in MSCs autophagy. Selective endoplasmic reticulum (ER) stress inhibitor salubrinal (SAL) was employed to determine whether PMP22 is mediated by ER stress. Our results demonstrated that VP1 played a promotive role in MSC autophagy. Overexpression of VP1 upregulated PMP22. PMP22 deficiency downregulated LC3B and thus inhibited autophagy. Furthermore, PMP22 expression was significantly suppressed by SAL. VP1 promotes MSC autophagy through upregulating ER stress-mediated PMP22 expression. VP1/ER stress/ PMP22 axis in autophagy may be a potential therapeutic target for EV71 infection-induced fatal neuronal damage.

## Introduction

Enterovirus 71, a single-stranded RNA virus, is one of the major causative pathogens of contagious hand, foot and mouth disease (HFMD)(1, 2) that mainly affects children under the age of 5. HFMD is an emerging public health issue worldwide, especially in Asia-Pacific countries(1, 3, 4). Although HFMD is commonly considered as a self-limited disease characterized by ulcerating vesicles in the mouth and viral rashes on hands and feet(5–7), a small proportion of cases are severe and even fatal due to cardiopulmonary or neurological complications(8, 9). EV71 infection accounts for at least 80% of severe cases and 90% of deaths in China according to the recent data (10). Increasing evidence indicates that EV71 may target human neurons in central nervous system (CNS), leading to neuronal degeneration and severe neurological disorders in fatal cases(11–13). Our previous study also showed that neuronal necrosis and neuronophagia were present in the brainstem in fatal EV71-infected cases(14). Despite these neurotropic characteristics of EV71 virus, the pathogenesis and molecular mechanisms of EV71-induced neuronal damage remain largely unknown.

Autophagy is an intracellular process that is mediated by a unique organelle named autophagosome and transports cytoplasmic components to the lysosomes for degradation(15, 16). The alteration of autophagy in the nervous system is associated with various neurodegenerative and neurometabolic disorders such as Alzheimer’s disease and Niemann-Pick disease(17–19). Autophagy can be observed using transmission electron microscopy (TEM) and can be assessed by measuring the conversion of microtubule-associated protein 1 chain 3 (LC3) to phosphatidylethanolamine (PE)-conjugated LC3 (LC3-II) localized in autophagosomal membranes, which reflects the number of autophagosomes or the degree of autophagy(20–22). EV71 has been shown to induce autophagy in infected human rhabdomyosarcoma and neuroblastoma cells(23, 24). Our previous study demonstrates that EV71 structural viral protein 1 (VP1) also induces autophagy in cultured primary EV71-infected brainstem neurons, which can be inhibited by endoplasmic reticulum (ER) stress inhibitor salubrinal (SAL)(25), suggesting an essential role of ER stress in VP1-induced autophagy.

ER stress is triggered by the accumulation of unfolded or misfolded proteins in ER(26, 27). Although the relationship between ER stress and autophagy is not yet fully understood, it is well established that there is a dynamic crosstalk between these two systems, and ER stress either stimulates or inhibits autophagy(26, 28, 29). Since ER stress and autophagy are commonly concurrent in some human pathologies, such as cardiovascular diseases, cancers, and neurodegenerative disorders(29–31), it is of great importance to identify ER stress-associated molecules as positive or negative regulators of autophagy. Peripheral myelin protein 22 (PMP22) is a transmembrane glycoprotein highly expressed in the myelinating Schwann cells of peripheral neurons, and majorly contributes to synthesis and function of myelin sheaths(32). In Schwann cells, newly synthesized PMP22 is transiently retained in ER and Golgi before transported to the plasma membrane(33, 34). Under pathological conditions, excessive mature or premature (unfolded or misfolded) PMP22 accumulates in ER and interacts with calnexin, a Ca^2+^-binding chaperone, leading to ER retention and activation of ER stress(35, 36). However, it remains unknown whether the relationship between PMP22 and ER stress is associated with autophagy.

In this study, we hypothesize that PMP22 is a downstream effector of ER stress and triggers activation of autophagy in response to EV71 capsid protein VP1. To confirm this hypothesis, we transfected mouse Schwann cells (MSCs) with VP1-expressing vectors to explore its effect on MSC autophagy and PMP22 expression. Our results showed that ER stress mediates the expression of PMP22 that is essential for MSC autophagy, suggesting an involvement of VP1/ER stress/PMP22 axis in the regulation of MSC autophagy. Targeting VP1/ER stress/PMP22 axis in autophagy may be a novel therapeutic strategy against EV71 infection-induced neuronal damage.

## Results

### Cloning and identification of VP1 cDNA

To determine if VP1 cDNA was successfully cloned into pEGFP-C3 vector, we prepared plasmids from transformed bacteria and digested them with *BamHI* and *XhoI*. The results of agarose electrophoresis showed that a band was located between 750 and 1000 bp following enzymatic digestion (Fig. 1), which is consistent with the size of VP1 cDNA (894 bp) based on the GenBank database. The sequencing results also indicated that the cloned fragment was identical to the VP1 cDNA sequence (supplementary Fig. 1), suggesting that VP1 cDNA was successfully cloned into the vector without any mutation.

**Figure 1.**
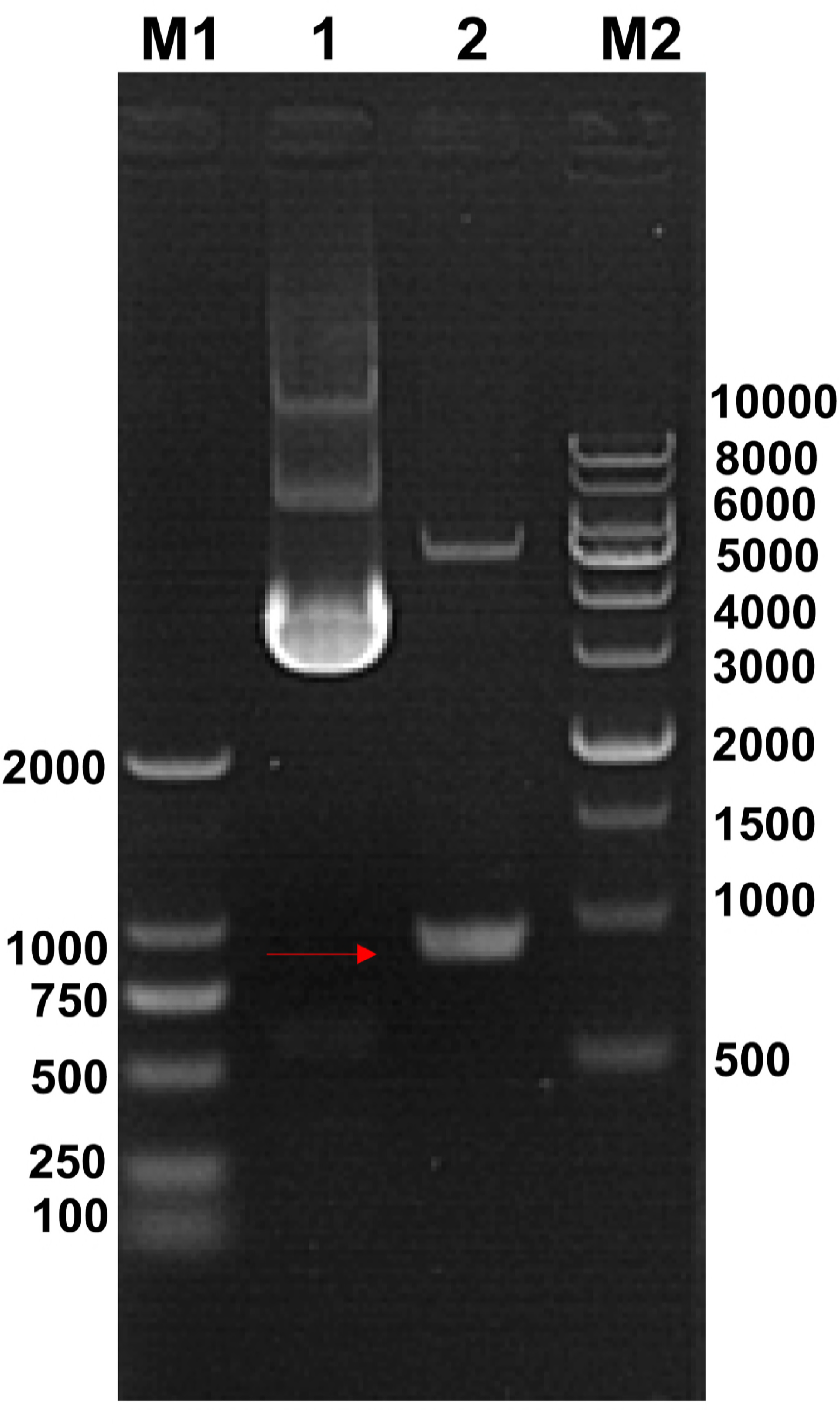
Cloning and identification of VP1 cDNA. Agarose electrophoresis for intact (lane 1) and restriction enzyme-digested (lane 2) VP1 cDNA cloning vector pEGFP-C3 plasmids. M: DNA marker.

### Overexpression of VP1 activates MSC autophagy

To examine whether VP1 has an effect on MSC autophagy, we analyzed the cellular and subcellular morphology of VP1-overexpressing MSCs using TEM. As shown in Fig. 2, VP1-overexpressing MSCs exhibited the features of autophagy such as swelling mitochondria, dilation and degranulation of rough ER, and vesicle-like dilation of Golgi(37), whereas the organelles in untransfected and GFP-transfected control MSCs were still morphologically normal. These results suggest that VP1 may activate autophagy in MSCs.

**Figure 2.**
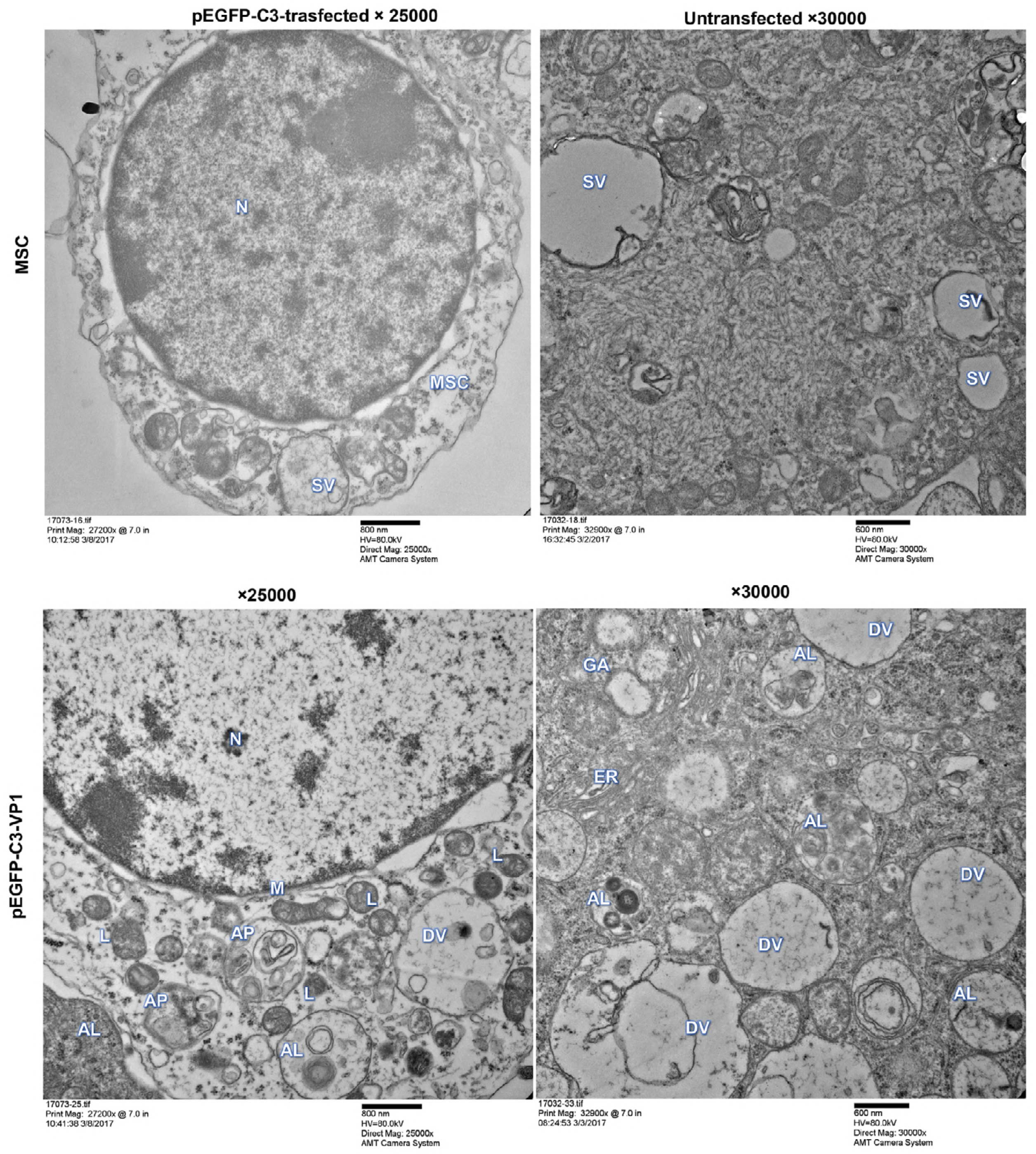
The effect of VP1 overexpression on MSC autophagy. MSCs were transfected with pEGFP-C3-VP1 plasmids for 48 h. Untransfected and pEGFP-C3-trasfected cells were used as blank and negative controls, respectively. Representative transmission electron microscopic images depict subcellular structures of MSCs. N: nucleus, M: mitochondrion, L: lysosome, AP: autophagosome, AL: autolysosome, DV: degradation vesicles, GA: Golgi apparatus, ER: endoplasmic reticulum, SV: secretory vesicles. MSC, mouse Schwann cell.

### Overexpression of VP1 markedly upregulates PMP22 expression in MSCs

Our previous study indicates an essential role of ER stress in VP1-induced autophagy in primary cultured EV71-infected brainstem neurons(14). In combination with the findings that PMP22 is abundant in Schwann cells and is closely associated with ER stress activation(32, 35, 36), we hypothesize that PMP22 might correlate with VP1 and play an important role in VP1-induced autophagy. To test this hypothesis, we detected the mRNA and protein expression of PMP22 in VP1-overexpressing MSCs. As shown in Fig. 3A, the mRNA expression of PMP22 was dramatically elevated in VP1-overexpressing MSCs compared with GFP-transfected cells. Immunofluorescent staining assay also showed similar results (Fig. 3B). These data indicate that VP1 is an upstream regulator of PMP22, suggesting a possible involvement of PMP22 in VP1-mediated activation of MSC autophagy.

**Figure 3.**
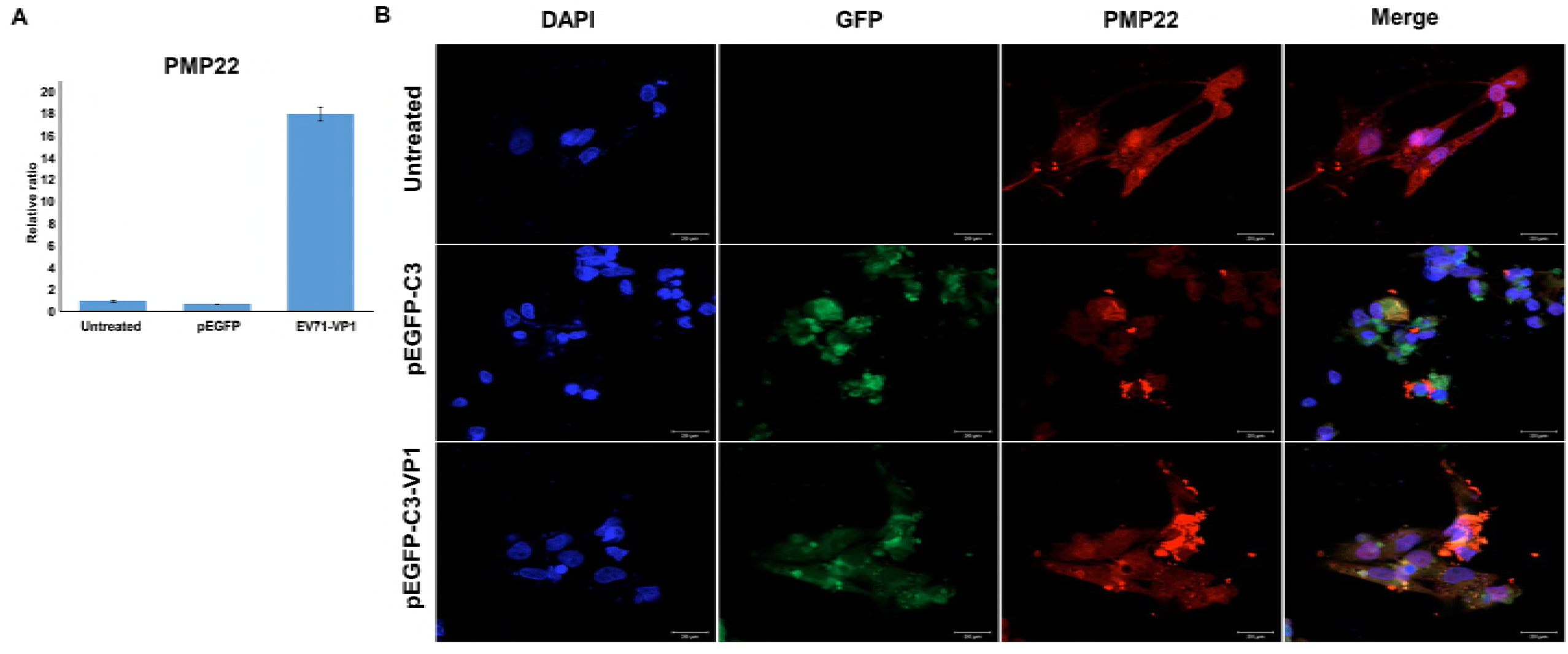
The effect of VP1 on PMP22 expression in MSCs. MSCs were transfected with pEGFP-C3-VP1 plasmids for 48 h. Untransfected and pEGFP-C3-trasfected cells were used as blank and negative controls, respectively. A. The mRNA expression of PMP22 was detected by qPCR. Data are expressed as the mean ± SE; \*\*\**P* < 0.001 vs. untransfected group; *n* = 3. B. Immunofluorescent staining for PMP22 in pEGFP-C3- or pEGFP-C3-VP1-transfected MSCs. GFP expression was used to monitor the transfection efficacy. Magnification: 400×. MSC, mouse Schwann cell; SE, standard error.

### PMP22 is essential for MSC autophagy

We next sought to investigate whether PMP22 is involved in MSC autophagy. PMP22 was knocked down by siRNA, which was confirmed by markedly decreased expression of PMP22 in siPMP22-transfected MSCs (Fig. 4A). Importantly, compared with siCtrl-transfected groups, knockdown of PMP22 significantly downregulated the expression of LC3 isoform LC3B-II, a gold standard autophagy marker (38), as shown in Fig. 4B and 4C. Consistently, the ratio of LC3B-II to LC3B-I in PMP22-deficient MSCs was also significantly lower than that in siCtrl-transfected groups (Fig. 4D). Furthermore, TEM images showed that there was no observable autophagic structure in siPMP22-transfected MSCs as compared with siCtrl-transfected cells (Fig. 5). Taken together, these data suggest that PMP22 is required for activation of autophagy in MSCs.

**Figure 4.**
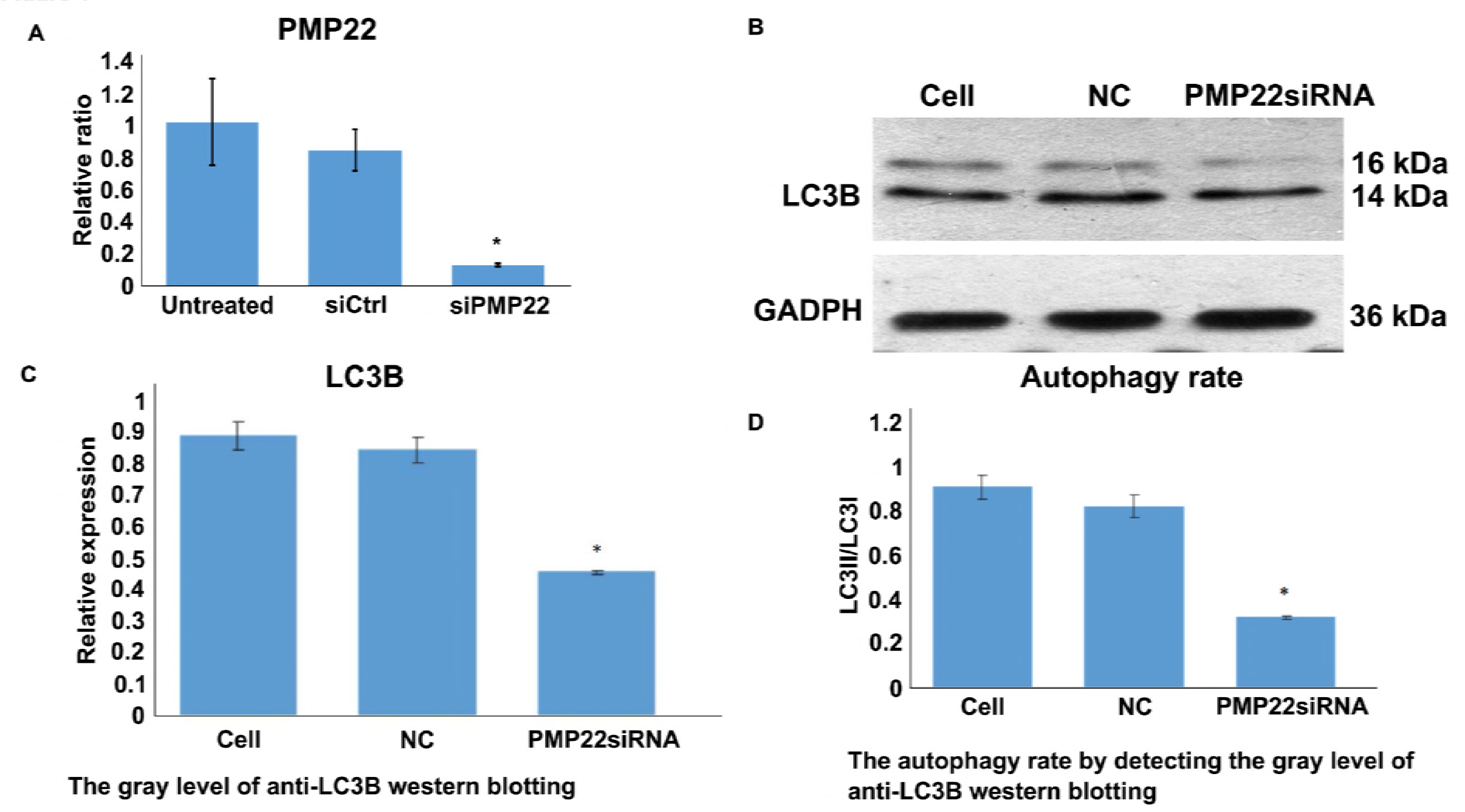
The effect of PMP22 knockdown on autophagy marker LC3B-II in MSCs. MSCs were transfected with siPMP22. Untransfected and scramble siRNA-trasfected cells were used as blank and negative controls, respectively. mRNA and proteins expression of LC3B were detected by real-time PCR (A) and Western blot assay (B), respectively. (C) Quantification of Western blot assay. (D) Ratio of LC3B-II to LC3B-I. Data are expressed as the mean ± SE; \*\*\**P* < 0.001 vs. untransfected group; n = 3. MSC, mouse Schwann cell; SE, standard error.

**Figure 5.**
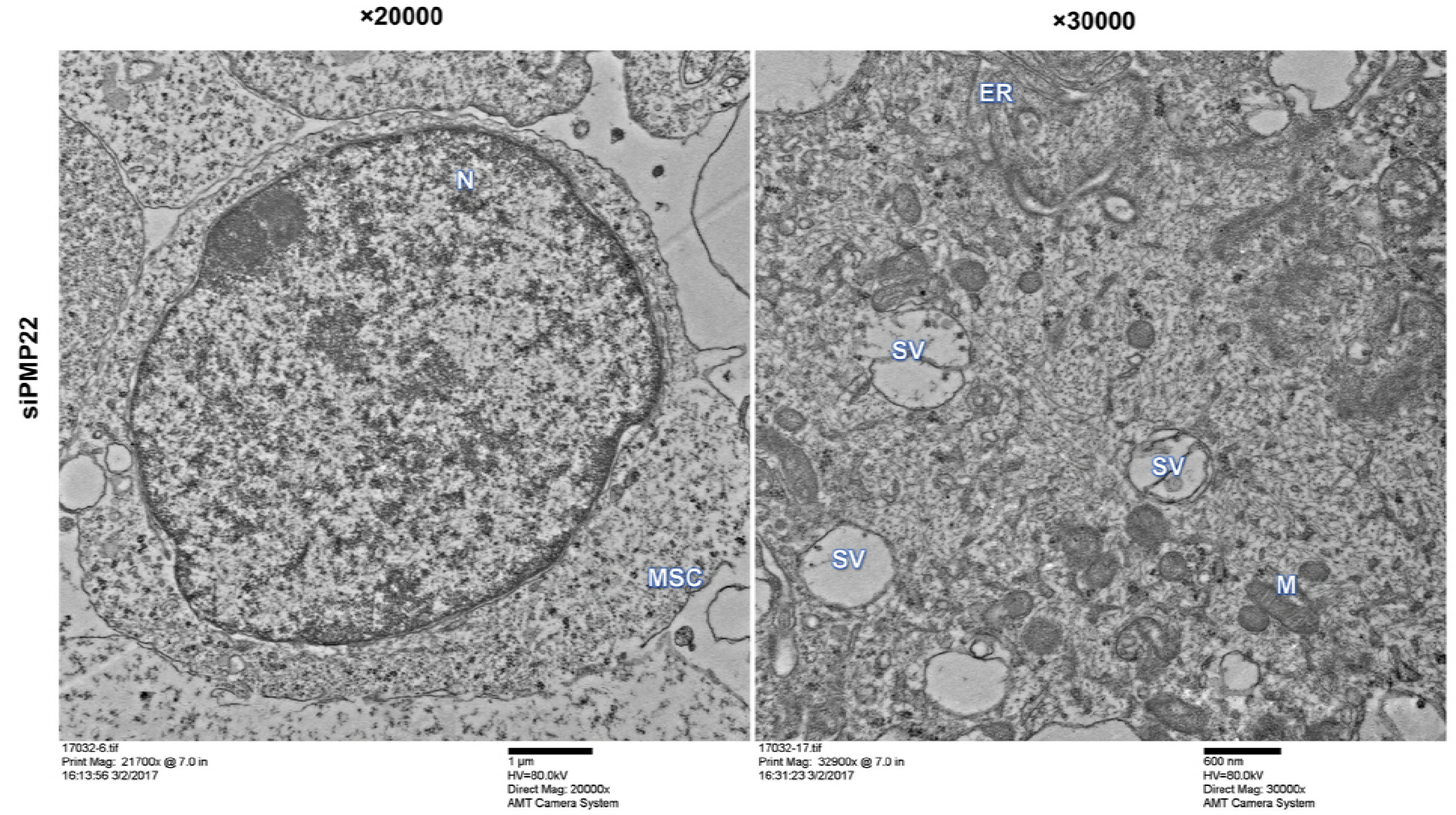
The effect of PMP22 knockdown on cellular and subcellular morphology of MSCs. MSCs were transfected with siPMP22 for 48 h. Untransfected and scramble siRNA-trasfected cells were used as blank and negative controls (siCtrl), respectively. Representative transmission electron microscopic images depict subcellular structures of MSCs. N: nucleus, M: mitochondrion, L: lysosome, AP: autophagosome, AL: autolysosome, DV: degradation vesicles, GA: Golgi apparatus. MSC, mouse Schwann cell.

### ER stress mediates PMP22 expression in MSCs

Since PMP22 is closely associated with ER stress and both PMP22 and ER stress are essential for activation of autophagy, we further sought to clarify the relationship between PMP22 and ER stress in MSCs using selective ER stress inhibitor SAL. As shown in Fig. 6A, compared with the control groups, mRNA expression of PMP22 was significantly downregulated following SAL treatment. Consistently, markedly weak fluorescent staining of PMP22 was also observed in SAL-treated MSCs (Fig. 6B), suggesting that PMP22 expression in MSCs is mediated by ER stress. These results indicate that VP1/ER stress/PMP22 signaling axis is an important component in MSC autophagy.

**Figure 6.**
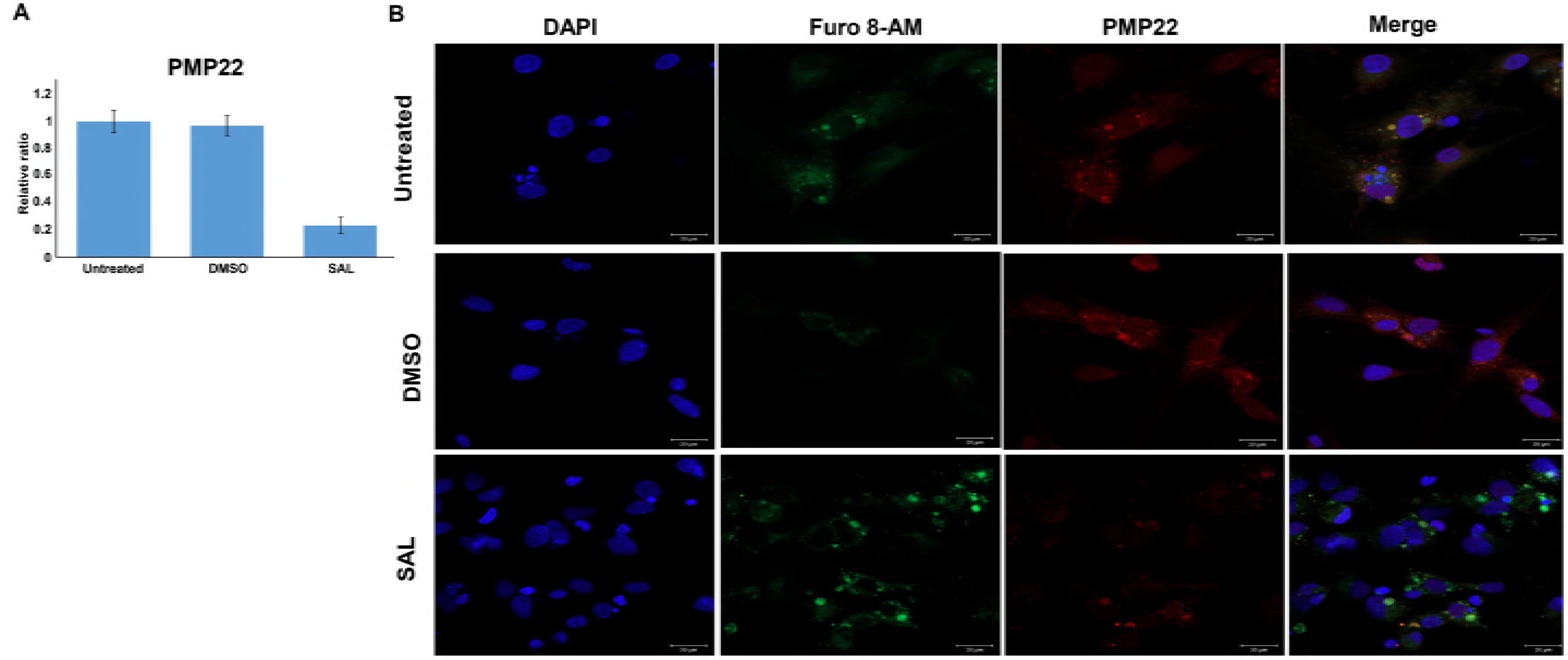
The effect of ER stress activation on PMP22 expression in MSCs. MSCs were treated with 15 μM of selective ER stress inhibitor salubrinal (SAL) for 48 h. Untreated and DMSO-treated cells were used as blank and negative controls. (A) The mRNA level of PMP22 was detected by real-time PCR. Data are expressed as the mean ± SEM; \*\*\**P* < 0.001 vs. untransfected group; n = 3. (B) Immunofluorescent staining (red) for PMP22 in DMSO- or SAL-treated MSCs. Furo 8-AM in SAL showed green fluorescence. Magnification: 400×. ER, endoplasmic reticulum; MSC, mouse Schwann cell; SE, standard error.

## Discussion

In the present study, we investigated the role and mechanism of EV71 capsid protein VP1 in MSC autophagy, and demonstrated for the first time that VP1 promotes MSC autophagy through ER stress-mediated PMP22 upregulation, suggesting VP1/ER stress/PMP22 axis as a novel potential target against EV71-induced neuronal disorder in severe HFMD cases.

EV71 possesses four structural proteins including VP1, VP2, VP3, and VP4. VP1 homodimers are the main component of the characteristic icosahedral capsid contributing to the pathogenicity and stability of EV71 virus to survive in the environment of the gastrointestinal tract(39, 40). In the present study, we demonstrated that VP1 plays a promotive role in MSC autophagy (Fig. 2), which is consistent with our previous findings(25). However, the effect of VP1-induced autophagy on MSC survival still remains unclear because autophagy plays dual roles in the nervous system. Excessive autophagy may be protective in chronic neurodegenerative diseases but detrimental in acute neural damages(18, 41). It has been reported that inhibition of EV71-induced autophagy in human rhabdomyosarcoma cells inhibits cell apoptosis at autophagosome formation stage and autophagy execution stage, but promotes apoptosis at the autophagosome-lysosome fusion stage. Furthermore, the inhibition of autophagy in the autophagsome formation stage or apoptosis decreases the release of EV71 viral particles, which is an effective strategy against virus infection(42). On the other hand, EV71-induced autophagy promotes viral replication in human rhabdomyosarcoma and neuroblastoma cells, and aggravates physiopathological parameters including weight loss, disease symptoms, and mortality in mouse models(23, 24). Further *in vitro* and *in vivo* studies are required to clarify the exact role of VP-induced autophagy in neuron cells.

In the present study, we also found that VP1 overexpression upregulated an important ER stress-associated protein PMP22(35, 36) in MSCs (Fig. 3), suggesting an involvement of ER stress activation in VP1-induced autophagy. It is well-established that excessive or premature PMP22 retaining in the ER induces ER stress(35, 36). However, the effect of ER stress activation on PMP22 expression hasn’t been investigated yet. Our data revealed for the first time that inhibition of ER stress significantly downregulated the expression of PMP22 in MSCs (Fig. 6), suggesting that PMP22 is a downstream effector of ER stress. It seems that there is a positive feedback loop between ER stress and PMP22 in MSCs. Furthermore, our results showed that, in PMP22-deficient MSCs, there was no morphological signs of autophagy and the autophagy marker LC3B-II was remarkably downregulated (Fig. 4 and 5), suggesting that PMP22 is essential for MSC autophagy. Interestingly, in an EV71-infected mouse model, VP1 was found co-localized with LC3 and/or autophagosome-like vesicles in neurons, and VP1 expression was positively correlated with LC3-II expression, aggregation and autophagosome formation(24). Upregulation of LC3-II expression was also observed in VP1-transfected HEK293 cells(25). Considering the regulatory role of VP1 in both MSC autophagy and PMP22 expression (Fig. 3), we conclude that VP1/ER stress/PMP22 pathway may play an important role in activation of MSC autophagy.

In summary, our data demonstrated that MSC autophagy can be activated by EV71 capsid protein VP1. Mechanistically, the expression of ER stress-associated protein PMP22 was significantly upregulated by VP1, suggesting that ER stress-mediated PMP22 upregulation is possibly responsible for VP1-induced autophagy activation. The VP1/ER stress/PMP22 axis may serve as a potential therapeutic target against EV71 infection.

## Materials and Methods

### Cell line and culture

Mouse Schwann cells (MSC) were purchased from ScienCell Research Laboratories (Carlsbad, CA, USA) and maintained in Schwann cell medium (ScienCell) containing penicillin (100U/mL)/ streptomycin (100 μg/mL) (Hyclone, Logan, UT, USA) in poly-L-lysine-coated (2 μg/cm^2^) flasks at 37°C in a humidified atmosphere of 5% CO_2_.

### Sample collection

EV71 were isolated from clinical specimens including throat, anal swabs and stools of HFMD patients with EV71 infection, and were provided by the Center for Disease Control and Prevention of Guangdong Province (Guangzhou, Guangdong, China). The patient was diagnosed by Guangxi Medical University (Nanning, Guangxi, China) based on the pathological analysis by Forensic Identification Center, Zhongshan School of Medicine, Sun Yat-sen University (Guangzhou, Guangdong, China).

### Gene cloning and transfection

Total RNA was extracted from EV71 using Trizol (Invitrogen, Carlsbad, CA, USA). The 894-bp VP1 cDNA was synthesized by reverse transcription polymerase chain reaction (RT-PCR) using the primer sets 5’-CCGCTCGAGGCCACCATGGGTGATGGAATTGCAGACATGA-3’ (forward) and 5’-CGCGGATCCTAGTGTTGTTATTTTGTCCCTACTTGTGC-3’ (reverse) (Genewiz, Suzhou, Jiangsu, China). The PCR products were then subcloned into pEGFP-C3 (Green Fluorescent Protein, GFP) expression vector (Clontech, Terra., USA) and sequencing was performed by Sangon Biotech (Shanghai, China). The results were compared with VP1 cDNA sequence reported by GenBank database. Cells were transiently transfected with plasmids using Lipofectamine 2000 (Invitrogen) following the manufacturer’s instruction.

### Small interfering RNA (siRNA)

siRNA against PMP22 (siPMP22) was from Santa Cruz Biotechnology (Dallas, TX, USA) and transfected using siRNA transfection reagent (Santa Cruz Biotechnology). Scramble siRNA (siCtrl) was used as a negative control.

### Quantitative real-time PCR (qPCR)

Total RNA was extracted from cells using Trizol (Invitrogen) following the manufacturer’s instructions and was reversely transcribed into cDNA using reverse transcriptase (Promega, Madison, WI, UDA). Real-time PCR was performed using SYBR Green qPCR SuperMix (Invitrogen) and the primers as shown in Table 1, following the manufacturer’s instruction. GAPDH was used as an internal control. Data were analyzed using the 2^−ΔΔCt^ method.

**Table 1.**
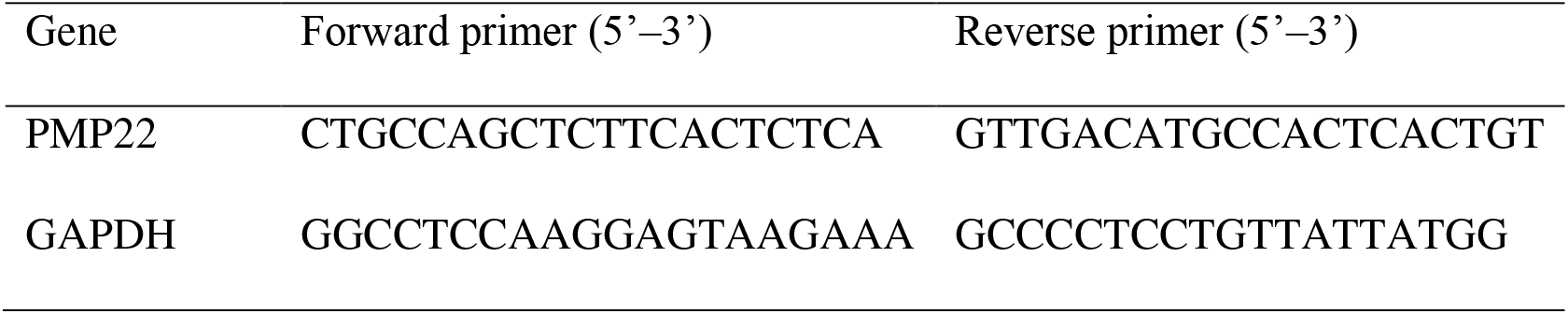
Real-time PCR primers.

### Western blot analysis

MSCs were lysed and the lysates were collected. Protein concentration was determined using BCA protein assay reagent (?). 50 ng of proteins were separated by 10% SDS-PAGE gel and transferred to polyvinylidene fluoride membranes. The membranes were then blocked with 5% nonfat milk powder in Tris-buffered saline and Tween 20 (TBST), and then incubated with anti-GAPDH (1:1000; Abcam, Cambridge, UK) or anti-LC3B (1:1000; Abcam) for 1–2 h at room temperature. Following 3 washes with cold TBST, the membranes were incubated with peroxidase-conjugated secondary antibody (1:4000; Thermo Fisher Scientific, Rockford, IL 61105 USA) for additional 1 h at room temperature. After 3 washes with TBST, the protein expression was detected using enhanced chemiluminescent development reagent (GE Healthcare, Little Chalfont, UK) and X-ray films.

### Immunofluorescence staining

MSCs were seeded on sterile coverslips 48 h after transfection and incubated overnight at 37 °C. Cells were then fixed with 4% paraformaldehyde for 30 min, followed by incubation with 0.2% Triton-X 100 at 4 °C for 5 min. After phosphate-buffered saline (PBS) washes, cells were blocked with 10% normal goat serum (Jackson ImmunoResearch, West Grove, PA, USA) for 30 min and incubated with anti-PMP22 antibody (Abcam) overnight at 4 °C. Cells were then incubated with fluorescence-conjugated secondary antibodies (Thermo Fisher Scientific, Waltham, MA, USA) for 1 h at room temperature. The images of stained cells were captured with a Leica camera (Leica, Wetzlar, Germany).

### Transmission electron microscopy (TEM) analysis

MSCs were prefixed with 2.5% glutaraldehyde for 2 h and postfixed with 1% osmic acid for additional 2 h at 4 °C, followed by gradient dehydration in 30%, 50%, and 70% ethanol (10 min each), 80%, 90%, and 95% acetone (10 min each), and 100% acetone (10 min twice). Cells were then embedded in the resin and stained with lead citrate. The stained cells were observed and imaged under a Hitachi H-7500 transmission electron microscope (Hitachi, Tokyo, Japan).

### Statistical analysis

All experiments were repeated at least three times. Data were expressed as the mean ± standard error (SE). Statistical significance was assessed using Student’s *t* test or one-way ANOVA with SPSS16.0 statistical software (SPSS Inc, IL, USA). *P* < 0.05 was considered statistically significant.

## Acknowledgements

None

